# SARS-CoV-2 RBD-Tetanus toxoid conjugate vaccine induces a strong neutralizing immunity in preclinical studies

**DOI:** 10.1101/2021.02.08.430146

**Authors:** Yury Valdes-Balbin, Darielys Santana-Mederos, Lauren Quintero, Sonsire Fernández, Laura Rodriguez, Belinda Sanchez Ramirez, Rocmira Perez, Claudia Acosta, Yanira Méndez, Manuel G. Ricardo, Tays Hernandez, Gretchen Bergado, Franciscary Pi, Annet Valdes, Tania Carmenate, Ubel Ramirez, Reinaldo Oliva, Jean-Pierre Soubal, Raine Garrido, Felix Cardoso, Mario Landys, Humberto Gonzalez, Mildrey Farinas, Juliet Enriquez, Enrique Noa, Anamary Suarez, Cheng Fang, Luis A. Espinosa, Yassel Ramos, Luis Javier González, Yanet Climent, Gertrudis Rojas, Ernesto Relova-Hernández, Yanelys Cabrera Infante, Sum Lai Losada, Tammy Boggiano, Eduardo Ojito, Kalet Leon Monzon, Fabrizio Chiodo, Françoise Paquet, Guang-Wu Chen, Daniel G. Rivera, Dagmar Garcia-Rivera, Vicente Verez-Bencomo

**Author notes:** These authors contributed equally (Yury Valdes-Balbin, Darielys Santana-Mederos, Sonsire Fernández, Belinda Sanchez Ramirez, Guang-Wu Chen, Daniel G. Rivera, Dagmar Garcia-Rivera, Vicente Verez-Bencomo).

## Abstract

Controlling the global COVID-19 pandemic depends, among other measures, on developing preventive vaccines at an unprecedented pace. Vaccines approved for use and those in development intend to use neutralizing antibodies to block viral sites binding to the host’s cellular receptors. Virus infection is mediated by the spike glycoprotein trimer on the virion surface via its receptor binding domain (RBD). Antibody response to this domain is an important outcome of the immunization and correlates well with viral neutralization. Here we show that macromolecular constructs with recombinant RBD conjugated to tetanus toxoid induce a potent immune response in laboratory animals. Some advantages of the immunization with the viral antigen coupled to tetanus toxoid have become evident such as predominant IgG immune response due to affinity maturation and long-term specific B-memory cells. This paper demonstrates that subunit conjugate vaccines can be an alternative for COVID-19, paving the way for other viral conjugate vaccines based on the use of small viral proteins involved in the infection process.

## Introduction

Control of SARS-CoV-2 infection focuses on development of preventive vaccines.^1^ Viral particles’ initial binding is mediated by the receptor binding domain (RBD) of the spike (S)-glycoprotein trimer to the host’s cell surface receptor, the angiotensin-converting enzyme 2 (ACE2).^2–5^ Most of the 200 COVID-19 vaccines in development^6^ aim to block this process.^1^ By focusing on the whole S-protein or its RBD as antigen, the primary goal is induction of anti-RBD antibodies that interfere with RBD-ACE2 interaction, blocking the first step of infection. Virus neutralization is mainly associated with antibodies against the receptor binding motif (RBM), a specific RBD region directly interacting with ACE2.^7^ This type of antibodies are not involved in antibody dependent enhancement (ADE).^8^

Key advantages of the well-known recombinant subunit vaccine platforms are their safety, stability at 2-8 ^o^C and facility to scale-up the production.^9^ While we should expect weak immunogenicity for such a small recombinant RBD protein (30 kDa), requiring repeated vaccination,^10^ it was found that recombinant RBD in alum is sufficient to induce a neutralizing immune response in laboratory animals,^11^ while its simplicity prompted subsequent evaluation in humans.^12^ However, most recombinant vaccines relied on large, RBD-containing macromolecular constructs as a way to increase the immunogenicity and on the use of potent adjuvants.^13^ In addition to the lower immunogenicity, small recombinant RBD exposes to the immune system not only the critical RBM surface but also RBD regions that are well-camouflaged at the virus surface. Antibodies directed to camouflaged RBD regions are not neutralizing, and therefore, an ideal construction should maximize the exposure of the RBM and not of such regions. We hypothesized that the orientation of RBD when conjugated to tetanus toxoid, exposes better the RBM surface increasing the level of neutralizing antibodies.^14–17^

The SARS-CoV-2 RBD comprises 193 amino acid residues from Thr333 to Pro527, including RBM 438-506 that interacts directly with ACE2. It contains eight cysteines forming four disulfide bridges, three of these stabilizing the RBD core and one within the RBM.^6^ Our recombinant RBD 319-541 was obtained in CHO-cells with intentionally extended sequence adding S-glycoprotein residues 527 through 541, in order to include an additional Cys538. This cysteine is usually connected to Cys590 in the S-glycoprotein. The extended sequence includes two N-glycosylation sites at residues Asn331 and Asn343 and two O-glycosylation sites at Thr323 and Ser325. The selected sequence results in an unpaired Cys538 intended to be used for chemical conjugation to the highly immunogenic carrier tetanus toxoid (TT). Here we report a promising vaccine candidate based on this high molecular weight conjugate with several copies of recombinant RBD per molecular unit. To our knowledge, chemically conjugated constructs and the immunogenic effect of conjugating viral proteins such as RBD to a protein carrier have not been assessed for SARS-CoV-2 or other coronaviruses. Here we demonstrate that the RBD-TT conjugate induces a potent immune response in laboratory animals, paving the way for their evaluation in human phase I and II clinical trials.^18^

### Construction of RBD-TT conjugates

Our design is based on the hypothesis that by conjugating several copies of the extended RBD to a large carrier protein, we can obtain a macromolecular construct displaying multivalent RBD. At the same time, the RBM will be well exposed (Fig. 1, represented in red) and more available for immune recognition.

**Fig. 1.**
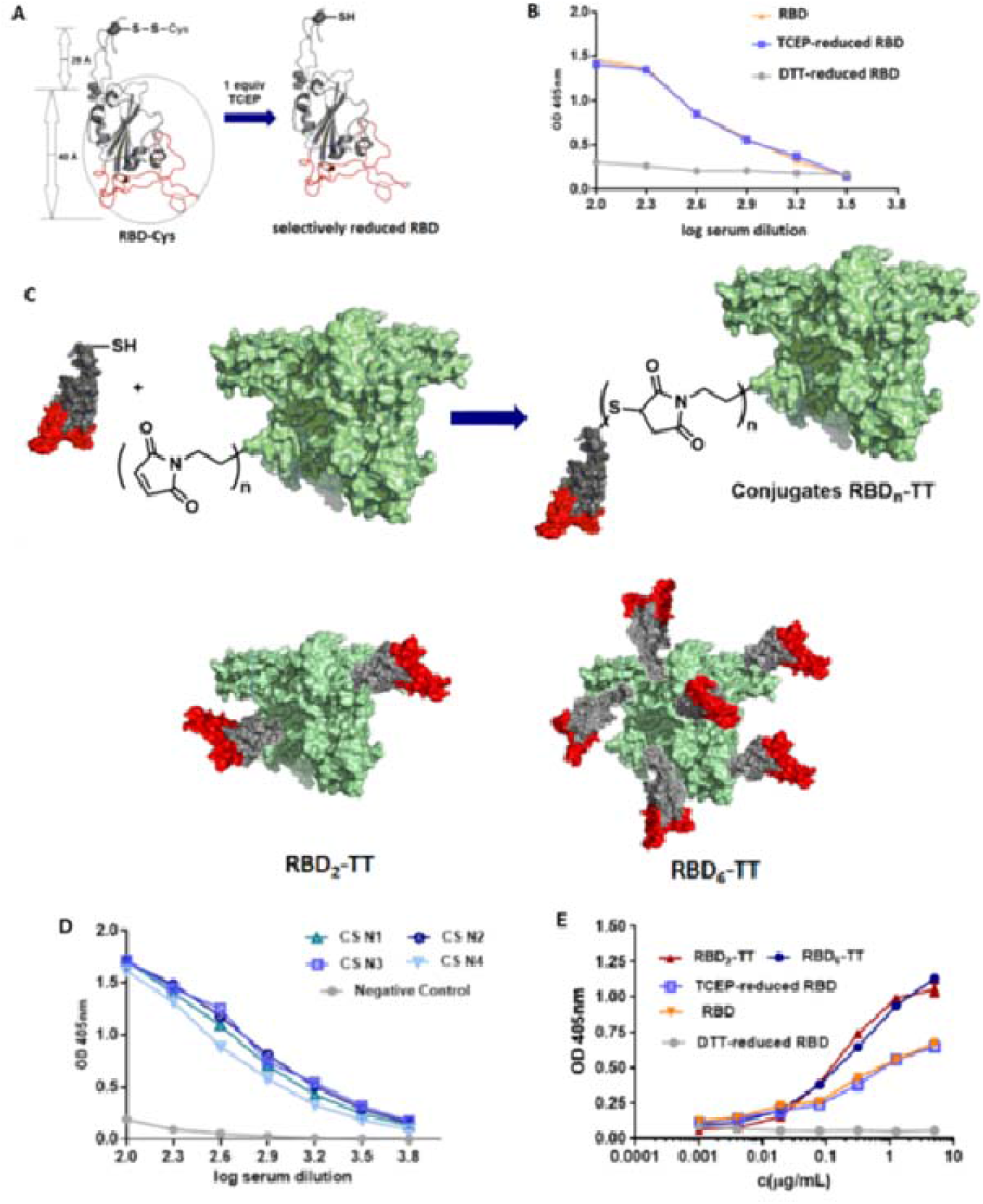
Synthesis of RBD-TT conjugates. (**A**) Reduction of RBD Cys538 using TCEP. (**B**) Recognition of RBD and reduced-RBDs by convalescent sera. (**C**) Conjugation of RBD with TT and representation of RBD_2_-TT and RBD_6_-TT. (**D**) Recognition of RBD-BSA conjugates by convalescent sera (CS), n=1-4. (**E).** Binding to ACE2 of RBD_2_-TT and RBD_6_-TT.

Inclusion of an additional free Cys538 in our extended RBD, while potentially useful for conjugation, could jeopardize extended-RBD folding, due to potential S-S rearrangement with the other 8 cysteines (scrambling). Nevertheless, we found that during fermentation and purification, Cys538 is spontaneously protected through an S-S Cys adduct with free cysteine present in the culture media. ESI-MS showed presence of the four native S-S bonds, indicating a correctly folded extended RBD (Fig. S2). Cysteinylated Cys538 was selectively reduced to free thiol with tris-(2-carboxyethyl)phosphine (TCEP)^19^ without affecting ACE2 recognition, while–for example–dithiothreitol (DTT) led to complete loss of its ACE2 binding capacity, suggesting loss of the antigen’s 3D structure (Fig. 1B).

To our knowledge, the immunogenic effect of TT as a carrier has not been assessed previously for SARS-CoV-2 or any other coronavirus. We have successfully used TT as a carrier protein for antibacterial carbohydrate-protein conjugate vaccines.^20,21^ The presence of multiple T- and B-cell epitopes of this highly immunogenic carrier^22^ might potentiate cellular immunity when compared to use of RBD alone. In addition, multimeric RBD-TT can simultaneously activate several B-cell receptors, thus enhancing B cell response.^23^

TT was activated with an average of 20–30 maleimide groups per mol of TT by reaction with *N*-succinimidyl 3-maleimidopropionate (SMP) followed by reaction with 2.5 or, alternatively, 10 equivalents of TCEP-reduced extended RBD, to produce conjugates bearing 2 or 6 mol, respectively, of RBD per mol of TT (Fig. 1c). The RBD_2_-TT and RBD_6_-TT conjugates were produced under good manufacturing practices (GMP) in 72% and 64% yield, respectively, and characterized by SE-HPLC and MS. Both conjugates recognize ACE2 slightly better than the original RBD, (Fig. 1E), confirming preservation of their structure and, probably, a better exposition of RBM. As convalescent serum usually contains TT antibodies, we prepared a RBD-bovine serum albumin (BSA) conjugate incorporating 6 RBD units per mol of BSA (RBD_6_-BSA), which was recognized well by various convalescent sera, proving conservation of the RBD antigenic properties after conjugation (Fig. 1D).

### Animal Immunogenicity

Immunization of BALB/c mice with the four different immunogens (Fig. 2) induces a strong IgG RBD–specific immune response as proven by ELISA. RBD_6_-TT/alum and RBD_2_-TT/alum were compared with RBD alone and with RBD_2_-TT without alum. After the first dose (T7 and T14, Fig. 2C) RBD_6_-TT/alum induces the highest level of anti-RBD antibodies. After the second dose all immunogens adsorbed in alum elicit better anti-RBD IgG levels than without alum (T21 and 28, Fig. 2C). The high and homogeneous early response for RBD_6_-TT/alum could be an important attribute for a vaccine in pandemic times. We explored early response to different dosage of RBD_6_-TT/alum, finding a dose-dependent response at day 7. At day 14, before the second dose, the response was very high even for the lowest dosage (T14, Fig. 2C).

**Fig. 2.**
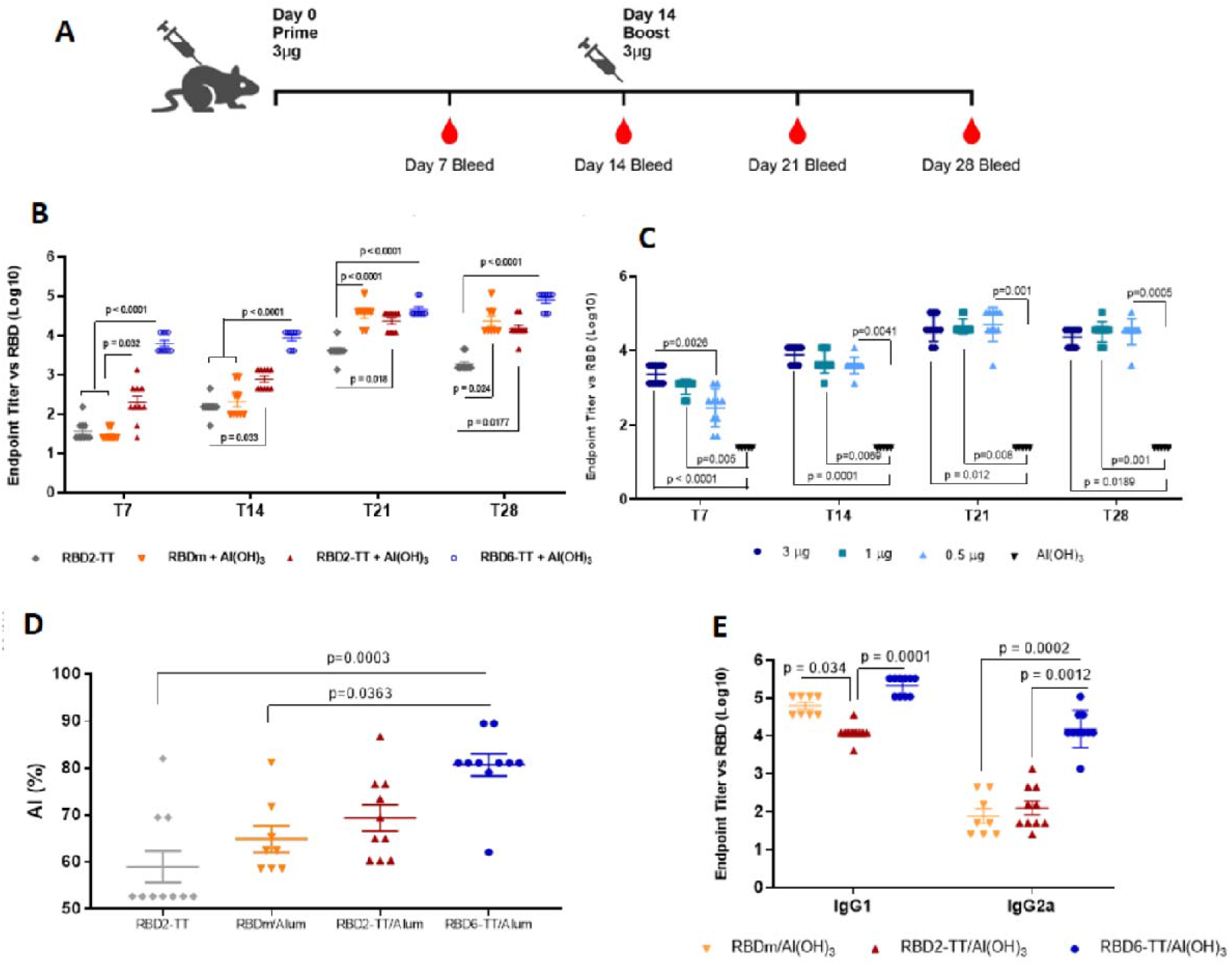
Immunization of BALB/c mice with RBD_2_-TT/ alum and RBD_6_-TT/alum compared to RBD and RBD_2_-TT. The serum of individual mice is represented by RBD/alum 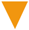, RBD_2_-TT/alum 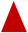, RBD_2_-TT 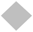, RBD_6_-TT/alum 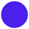 (**A**) Immunization protocol. (**B**) anti-RBD–specific IgG at days 7, 14, 21, and 28. (**C**) Dose response to RBD_6_-TT/alum at days 7 and 14. (**D**) Avidity index of antibodies elicited at T28. (**E**) RBD-specific IgG1 and IgG2a.

To evaluate possible immunological advantages of the RBD-TT conjugate, we studied affinity maturation, of antibodies elicited by RBD_6_-TT compared with the rest of immunogens. There was an increase in the avidity index (AI).^24^ The highest value of 81% for antibodies induced by RBD_6_-TT is consistent with a more pronounced affinity maturation. (Fig. 2D). The Th1/Th2 balance can be modulated by vaccination and was also evaluated (Fig. 2E). A biased Th2 immune response was observed for RBD_2_-TT/alum (IgG2a/IgG1 ratio 0.54) and RBD/alum (IgG2a/IgG1 ratio 0.40), while RBD_6_-TT/alum displayed more balanced Th1/Th2 immunity (IgG2a/IgG1 ratio 0.81).

Fig. 3 shows the induction of memory antigen-specific B and T-cells, an important property of conjugate vaccines. Mice immunized with conjugate RBD_6_-TT/alum were compared to mice receiving RBD/alum. Both groups developed a primary immunity as shown previously (Fig. 2B). After two doses at T28, splenocytes purified from both groups were intravenously transferred to naïve mice that were then boosted by a single dose of 3 μg RBD/alum (Fig. 3B). Mice receiving splenocytes from RBD_6_-TT/alum responded with a strong secondary RBD–specific IgG response (titer 10^3^-10^4^), while those receiving splenocytes from RBD/alum did not (results not shown). This finding demonstrated the presence of RBD-specific memory B cells in transferred splenocytes, which were able to be activated in the presence of RBD/alum (alternative SARS-CoV-2 virus) stimuli.

**Fig. 3.**
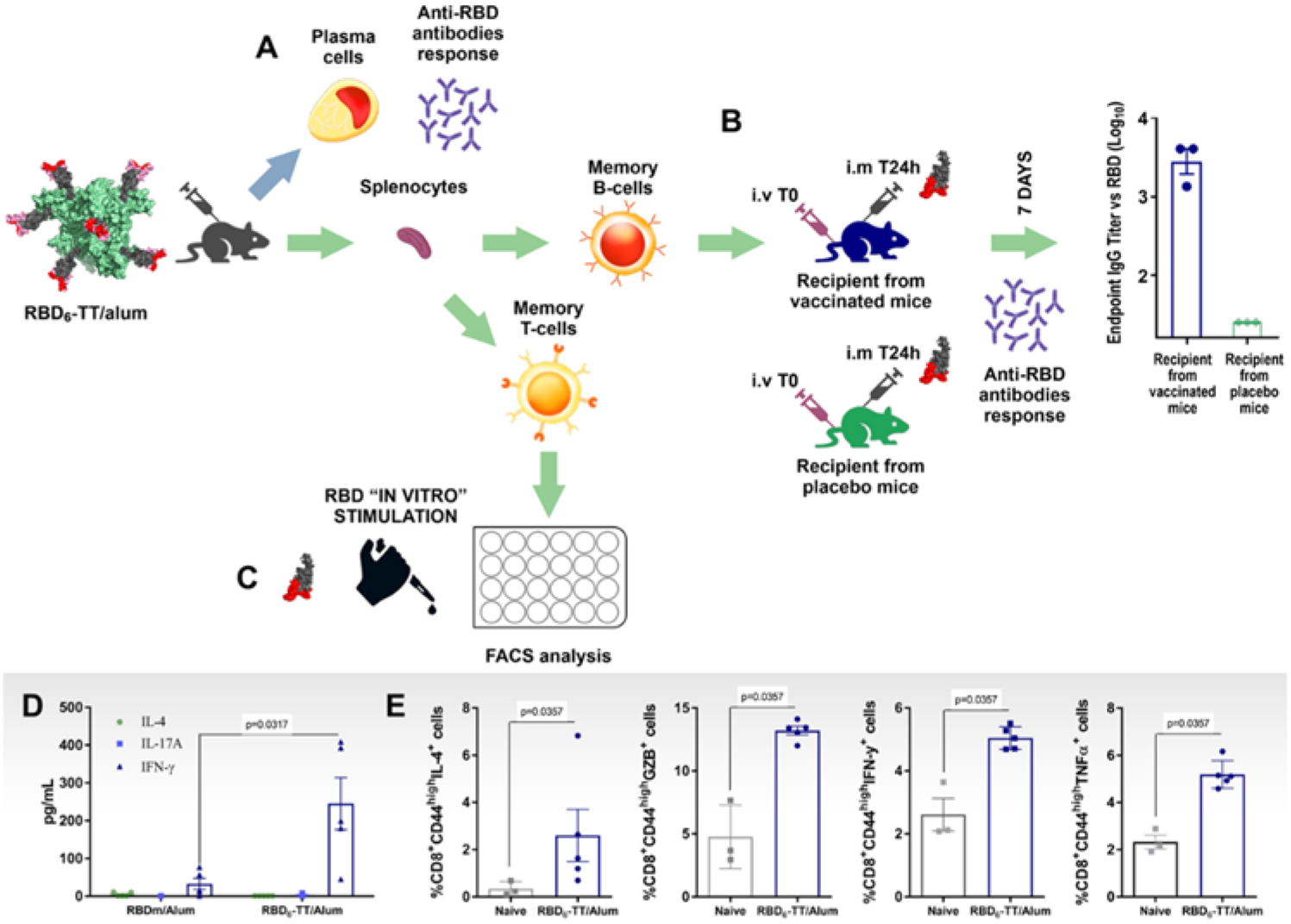
Memory B and T cells induced by RBD_6_-TT. (**A**) Primary immune response to RBD_6_-TT/alum (green arrows). (**B**) Classical passive transfer of splenocytes from RBD_6_-TT/alum BALB/c and stimulated with RBD/alum (strong secondary response after day 7). (**C**) T-cell stimulation with RBD. (**D**) Cytokine secretion after *in vitro* RBD stimulation (**E).** % RBD-specific memory T CD_8_^+^CD_44_^high^IL-4^+^; % RBD-specific memory T CD_8_^+^CD_44_^high^Granzyme^+^; % RBD-specific memory T CD_8_^+^CD_44_^high^rFNβ^+^; % RBD-specific memory T CD_8_^+^CD_44_^high^TNF_α_^+^

Specific CD8+ T cells also play an important role in protection, as recently demonstrated.^25^ To evaluate the specific T-cell response, we compared RBD_6_-TT/alum and RBD/alum immunized mice. After *in vitro* RBD stimuli, splenocytes from mice immunized with RBD_6_-TT secreted higher levels of IFNγ compared to those immunized with RBD/alum (Fig. 3D), suggesting a Th1 pattern, while IL-4 (characteristic of Th2 pattern) and IL-17A (characteristic of Th17 pattern) were not detected. Frequency of CD8^+^CD44^high^ memory T-lymphocytes producing IFN-γ, TNF-α and Granzyme B increased significantly in RBD_6_-TT immunized mice with respect to control mice (Fig. 3E) as shown by flow cytometry, indicating the relevant activation of cytotoxic T immune memory.

### Antibody functionality

We evaluated antibodies’ ability to block interaction between the virus and its receptor,^10^ using the molecular Virus Neutralization Test (mVNT50)^22^ and the conventional Virus Neutralization Test (cVNT50).^23^ mVNT50 evaluates inhibition of interaction between recombinant RBD and ACE2 at the molecular level; at the cellular level, cVNT50 evaluates inhibition of interaction between the live virus and Vero E6 cells bearing ACE2 receptors. Antibodies resulting from immunization of Balb/c mice with two doses of RBD_2_-TT/alum and RBD_6_-TT/alum were compared to antibodies elicited after immunization with RBD/alum. mVNT50 showed a high level of inhibition for all sera (Fig. 4a), indicating that all tested antibodies displayed a similar efficacy in interfering with RBD-ACE2 interaction at the molecular level. cVNT50 (Fig. 4B) showed sharp differences between sera from animals immunized with RBD/alum and those with both conjugates. For RBD/alum, the neutralization titer was 232; for both conjugates, there was a higher level of virus neutralization: 1303 for RBD_2_-TT and 2568 for RBD_6_-TT. The mVNT50/cVNT50 ratio was 0.143, 0.732 and 1.08 for RBD, RBD2-TT and RBD6-TT respectively. While antibodies neutralizing the virus are mainly directed at the RBM,^7^ there are antibodies recognizing soluble RBD not only by the RBM but also on a different region, as shown in Fig. 4C. This type of “lateral” antibodies could interfere in mVNT50 with soluble RBD, but probably will not recognize this RBD region that is camouflaged at the virus surface.

**Fig. 4.**
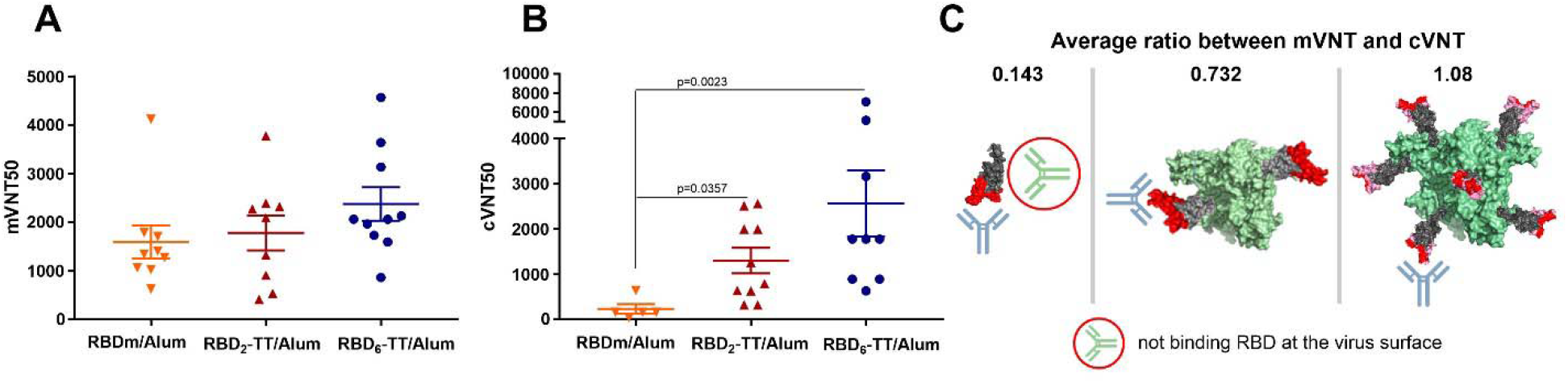
Virus neutralization by RBD antibodies induced by conjugates. (**A**) mVNT_50_ representing the serum dilution giving 50% inhibition ACE2-RBD interaction. (**B**) cVNT_50_ measured as serum dilution giving 50% of virus neutralization (**C**) mVNT50/cVNT50 ratio and a possible schematic representation of the differences found.

Based on the results presented here, GMP batches of the conjugates RBD_2_-TT and RBD_6_-TT were obtained and absorbed on alum as final vaccine candidates. A phase I clinical trial^24^ was initiated in October 2020, and after preliminary results confirming a better performance of a vaccine based on RBD_6_-TT/alum, this advanced on December 2020 to a phase II clinical trial with 910 subjects.^25^ The encouraging interim results of all those trial envisage moving forward to a phase III trial in March 2021. The resulting vaccine, the first one developed and produced in a Latin-American country, has important advantages to be considered in the future: a) it induces a strong IgG neutralizing antibody as well as specific T-cell response, b) the well-known safety record of this vaccine platform is further confirmed during clinical trial, c) the storage and distribution temperature of 2-8 ^o^C is the conventional for other vaccines and suitable for a rapid distribution worldwide, and finally, d) it can be adapted to existing vaccine production capacities available at several countries. We hope this vaccine can contribute to win the battle against the SARS-CoV-2 pandemic.

## Supporting information

manuscrito v1-03-2021.docx

## Acknowledgments

We thank Rolando Pérez, Luis Herrera, Agustin Lage and Eduardo Martinez (BioCubaFarma), for advice and support to the project, Lila Castellanos and Gail Reed for editing English version; as well as the Fondo de Ciencia e Innovacion (FONCI) for financial support (Project-2020-20). We are grateful to the Leibniz Institute of Plant Biochemistry, Germany, for support in structural characterization.

## Author contributions

Y.V.B., D.G.R., and V.V.B. designed and lead the study. S.F.is the manager of the project. D.S.M. and D.G.R. designed conjugation protocols and L.Q., U.R., J.P.S,. Y.M., H.G., and M.G.R performed chemical conjugation. L.R., B.S.R., R.P., C.A.,T.H., G.B., F.Pi., A.V., performed the immunologic assays, M.F. and R.O. led the clinical care of the animals, F. C., R.G., M.L. led the analytical chemistry of the conjugates and vaccines J.E., N.G., and A.S. performed the virologic assays L.A.E., Y.R., and L.J.G. performed mass-spectra studies of the conjugates G.R., E.R-H., Y.C.I., S-L.L.,T.B., E.O., K.L.M., C,F, and G.W.C. led the CHO-cells RBD preparation, Y.C., F.C. and F.P. supported and contributed to the study design and analysis. All authors revised the manuscript and approved submission.

## Supplementary material

**Material and Methods**

**Table I-II**

**Fig. S1-S9**

