## Supplementary material for "SARS-CoV-2 RBD-Tetanus toxoid conjugate vaccine induces a strong neutralizing immunity in preclinical studies": manuscrito v1-03-2021.docx

*These authors contributed equally.

**Materials and Methods**

**Analysis.** The HPLC system was equipped with a Knauer/Azura quaternary pump system P 6.1L and Knauer a K-2501 UV/Vis Detector. Data acquisition and analysis were performed with ClarityChrom (DataApex) software. The analyses were performed by high performance size exclusion with PBS pH 7 buffer on a Superdex 75 Increase® 10/300 GL column and Superdex 200 Increase® 5/150 GL column (GE Healthcare) at a flow rate of 0.8 ml/min and 0.25 ml/min, respectively. SDS-polyacrylamide gel electrophoresis (SDS-PAGE) was performed in a gradient gel (4-20) % acrylamide, prepared with 10 µg for conjugates *per lane* and 5 µg for tetanus toxoid and RBD. The gel was stained with Coomassie blue R250 and unstained with a methanol/acetic acid/water solution. Final analysis was performed with Bio-rad GS-800 densitometer and Quantity One software. Figures showing 3D structures of RBD and TT proteins were performed using the PyMOL Molecular Graphics System from PDBs 6M0J and 5N0B, respectively.[1]

[1] De Lano W.L., Pymol. South San Francisco, CA: De Lano Scientific **2002**

**Mass Spectrometry analysis.** A volume equivalent to five micrograms of the protein solution treated with *N*-ethyl maleimide (NEM), and deglycosylated with PNGase-F was desalted by using ZipTips C18 (Millipore). The desalted protein was loaded into the metal coated nanocapillary for ESI-MS analysis. The mixture of tryptic peptides and the deglycosylated RBD was analyzed in a hybrid orthogonal QTof-2$^{TM}$ tandem mass spectrometer (Micromass, Manchester, UK) by a spraying the sample into ion source using 1200 y 35 volts for the capillary and the entrance cone, respectively. The ESI-MS were acquired from *m/z* 200-2000 and the multiply-charged ions were manually fragmented by collision induced dissociation using appropriated collision energies (20-50 eV) to obtain structural information in the MS/MS spectra. The ESI-MS/MS of tryptic peptides with $z \geq 3+$ were deconvoluted by MaxEnt 3.0. Argon was used as a collision gas. The multiply-charged ESI-MS spectrum (*m/z* 400-3000) of the protein deglycosylated with PNGase F was deconvoluted (mass 3000-70000) by using MaxtEnt1.0 software. The theoretical *m/z* for tryptic peptides as well as for the intact protein was calculated by using the peptide and protein editor available in the MassLynx v4.1 software (Micromass, UK).

**Production of monomeric RBD (Arg319-Phe541-(His)$_6$) recombinant protein:** The coding sequence for RBD319-541 fused to a C-terminal hexahistidine tag was optimized for mammalian cell expression (hamster, *Crycetulus grisesus*), using the online gene optimization tools provided by Eurofins (Germany). The resulting nucleotide sequence was assembled and amplified by PCR using gene fragments synthesized by Eurofins and oligonucleotides synthesized by CIGB (Cuba) and cloned into an intermediate vector containing CMV promoter and mouse Ig $V_H$ signal sequence gene. The whole expression cassette was subsequently re-cloned into the lentiviral vector pL6WBlast, kindly provided by CIGB. HEK-293T cells were co-transfected with the lentiviral vector containing the gene of interest plus the mixture of auxiliary plasmids pLPI, pLPII and pLP/VSV-G, and used to produce lentiviral particles. CHO-K1 host cells were transduced with lentiviral particles and grown in 96 well plates in the presence of the selection drug blasticidine. Supernantants were screened for secretion of recombinant RBD by ELISA, and transduced cells showing the highest secretion levels were adapted to grow in suspension in serum-free medium (a mixture of PFHMII with a CIM proprietary medium) with shaking. Recombinant RBD319-541 was purified from cell culture supernatant by immobilized metal affinity chromatography (IMAC) with Ni-NTA Sepharose. Monomeric RBD was then isolated by size exclusion chromatography (SEC) using Superdex 200®.

**ESI-MS analysis of recombinant RBD:** Recombinant purified RBD ($Arg^{319}$-$Phe^{541}$-$(His)_6$) was deglycosylated with PNGase-F and treated with NEM before electrospray ionization mass spectrometry analysis (ESI-MS). The ESI-MS showing the multiply-charged ions is shown in Fig. S1A. This mass spectrum was deconvoluted and the experimental molecular mass was found to be 26981.82 Da (Fig. S1B), which is approximately 119 Da higher than the expected (26 982.22 Da), indicating that the recombinant protein is cysteinylated. Also it is appreciable the heterogeneity introduced by the O-glycosylation showing the presence of two O-glycoforms (HexNAc+Hex+$SA_2$ and HexNAc+Hex+SA). The assignments for all signal detected in Figure S1B are summarized in Table I. The experimental molecular masses agreed very well with the expected values considering the four disulfide bonds, the two potential N-glycosylation sites transformed into Asp residues during the PNGase-F treatment and $Cys^{538}$ cysteinylated. It was also observed that NEM was added partially at the N-terminal residue of the protein.

To confirm this, the protein was digested with trypsin by using an in-solution buffer-free digestion protocol and analyzed by ESI-MS.[2] The ESI-MS spectrum of the proteolytic peptides is shown in Figure 2A and Table II summarizes the corresponding assignments. The four disulfide bonds [($C^{336}$-$C^{361}$), ($C^{379}$-$C^{432}$), ($C^{391}$-$C^{525}$) and ($C^{480}$-$C^{488}$)] present in the spike protein of SARS-CoV-2 were detected. The signal detected at *m/z* 475.20 (3+) (Figure 1C, left panel) was assigned to the C-terminal peptide $C^{538}$VNFHHHHHH$^{547}$ containing cysteinylated $Cys^{538}$ (see Table S1). The MS/MS spectrum confirmed this assignment. In the same expanded region (*m/z* 468-483), a small signal at *m/z* 477.19 (3+) indicated a minority presence of the C-terminal peptide $C^{538}$VNFHHHHHH$^{547}$ modified with NEM at $Cys^{538}$, implying that only a small portion of the $Cys^{538}$ thiol group was free while the majority was cysteinylated prior to the addition of NEM.

**Reduction of RBD-Cys to obtain the RBD with free Cys538-thiol:** 14 mL of a solution of recombinant RBD (12 mg/mL, 5.6 μmol based on the MW of the monomer) in 35 mM PBS pH 7.4 and 0.5 mM EDTA was treated with an aliquot (1.6 mL) of a freshly prepared solution of tris(2-

[2] Betancourt, L. H. Espinosa, L. A.; Ramos, Y., Bequet-Romero, M., Rodríguez, E. N., Sánchez, A., Marko-Varga, G., González, L. J., Besada, V. *Targeting hydrophilic regions of recombinant proteins by MS using in-solution buffer-free trypsin digestion. Eur. J. Mass Spectrometry* **2020**, *26*, 230-237. doi: 10.1177/1469066719893492.

carboxyethyl)phosphine (TCEP)[3] hydrochloride (1 mg/mL, 5.6 μmol, 1 equiv). The reaction was incubated for 10 min at room temperature, time after which the conversion to the decysteinylated RBD was confirmed by ESI-MS analysis. For this, a small sample was treated with NEM (5 mM), deglycosylated and digested with trypsin as described above. The deprotection of thiol group at Cys538 was demonstrated by tracking the $(Cys538+Cys)^{3+}$signal previously detected at *m/z* 475.19, which disappeared and was transformed into a signal at *m/z* 477.19 (Fig. 3C, right panel), assigned to $(Cys538+NEM)^{3+}$. The other four disulfide bonds were not considerably affected in the treatment with TCEP, as they were detected with similar intensities. The TCEP-reduced RBD was also analyzed by SDS-PAGE under non-reducing conditions in comparison with DTT-reduced and non-modified RBDs (Fig. S3). Reduction with DTT led to a significant loss of the RBD three-dimensional structure, while TCEP-reduced and non-modified RBD (i.e., recombinant RBD-Cys) showed a similar behavior.

**Activation of tetanus toxoid with maleimide:** 10 mL of a solution of tetanus toxoid (TT, 5 mg/mL, 50 mg, MW 150 kDa, 0.33 μmol) in 100 mM 4-(2-hydroxyethyl)piperazine-1-ethanesulfonic acid (HEPES) pH 7.8 was treated by slow addition with 187 μL of a freshly prepared solution of *N*-succinimidyl 3-maleimidopropionate (SMP) in DMSO (75 mg/mL, 53 μmol, 160 equiv of SMP per mol of TT). The mixture was gently stirred for 1 h at room temperature and then purified by diafiltration (100 kDa cut-off) with 35 mM PBS pH 7.4 and 5 mM EDTA; and finally concentrated to 18 mg/mL, as measured by Lowry's method.[4] The stoichiometry of maleimide linkers covalently attached to TT was determined to be 20-30, as measured by a reversed Ellman method.[5]

**Conjugation of RBD to maleimide-activated TT:** 13.5 mL of a solution of RBD with free Cys538-thiol (12 mg/mL, 5.4 μmol), arising from the TCEP reduction procedure, was treated with a solution of maleimide-activated TT (4.5 mL, 18 mg/mL, 0.54 μmol) in 35 mM PBS pH 7.4 and 5 mM EDTA (final concentration RBD is 9 mg/mL and activated-TT is 4.5 mg/mL, molar ratio

[3] Daniel J. Cline, Sarah E. Redding, Stephen G. Brohawn, James N. Psathas, Joel P. Schneider, and Colin Thorpe. *New Water-Soluble Phosphines as Reductants of Peptide and Protein Disulfide Bonds: Reactivity and Membrane Permeability. Biochemistry* 2004, 43, 15195-15203.

[4] Lowry, O. H., N. J. Rosebrough, A. L. Farr, and R. J. Randall. *Protein measurement with the Folin phenol reagent.* (1951) *J. Biol. Chem.* 193:265–275.

[5] Ellman, G. L. *A colorimetric method for determining low concentrations of mercaptans*. (1958) *Arch. Biochem. Biophys.* 74:443–50.

RBD/TT 10:1). The reaction mixture was gently stirred for 16 h at 5 ± 3 °C and then was treated with 0.6 mL of a cysteamine hydrochloride solution (5 mg/ml) and incubated for 30 min. The decrease in the peak area for RBD was used as a parameter to monitor the conjugation progress and to quantify the amount of RBD incorporated into the RBD-TT conjugate. Thus, the average stoichiometry of RBD linked to TT was found to be 6.4 ($RBD_6$-TT). Conjugate $RBD_6$-TT was obtained in 64% yield (based on RBD) after purification by diafiltration (100 kDa cut-off) with 20 washes of 1 mM PB pH 7.0. Using the same conjugation procedure but with a molar ratio RBD/maleimide-activated TT of 2.5:1, a conjugate was obtained in 72% yield showing an average stoichiometry of RBD linked to TT of 2 ($RBD_2$-TT). A $RBD_6$-BSA conjugate was also obtained in 92% from maleimide-activated BSA using the same protocol. The purified RBD-TT and RBD-BSA conjugates were characterized by SE-HPLC, SDS-PAGE, Western blot and ELISA.

**ELISA assays**

**Recognition of human ACE2:** Microtiter plates (High binding, Costar, USA) were coated with 50µL/well of ACE2-mFc (5 µg/mL) in carbonate-bicarbonate buffer, 0.1 M (pH 9.6) and incubated overnight at 4 °C. Plates were blocked with 200µL/well of 2% of Non Fat Dry Milk (NFDM) in PBS-Tween 20 0.05% (PBST) for 1 h at 37 °C. Serial dilutions of the RBD-TT conjugates were prepared in 0.2 % NFDM/PBST and 50µL were added and incubated for 2 h at 37 °C. RBD and 6×His tagged PD-L1 were used as positive and negative control, respectively. The bound protein was detected using 50 µL/well of RBD specific rabbit polyclonal antibodies (PAbs, 100µg/mL) for 1 h at 37°C. Next, 50 µL/well of a peroxidase conjugated anti-rabbit IgG monoclonal antibody (MAb, 1:10 000) was added and incubated for 1 h at 37 °C. The reaction was visualized by addition of substrate 3,3’,5,5’-Tetramethylbenzidine (TMB) (BDBiosciences) and stopped with $H_2SO_4$ (1 M). The absorbance at 450nm was measured using a microwell system reader (Organon Teknica, Salzburg, Austria). All incubations were followed by three washing steps with PBST.

**Animal experiments**

Immunogenicity evaluation of each conjugate was performed in BALB/c and C57BL/6 mice (6-8 weeks of age, 15-20 g), New Zealand rabbits (7-8 weeks of age, 1.5 -1.8 kg) and Syrian hamsters (7-8 weeks of age, 35-45 g). Also, elder C57BL/6 mice (66-68 weeks of age) were used. All animals were supplied by the National Center for Laboratory Animals Breeding (CENPALAB),

Havana, Cuba with their health certificates. All protocols were approved by the Finlay Vaccine Institute Ethical Committee.

Independently of the animal model, all schemes consisted of two intramuscular doses, 14 days apart, and sera were collected previous to immunization at 7, 14, 21 and 28 days after, for RBD-specific antibodies kinetics.

Six groups of 10 mice were injected with: a) 1 or 3 µg/100 µL of RBD-TT conjugates adjuvated or not with 500 µg of $Al(OH)_3$; b) 3 µg/100 µl of RBD adjuvated with 500 µg of $Al(OH)_3$; c) 500 µg/100 µL of $Al(OH)_3$ as control. Cytokine and T memory response was evaluated in BALB/c mice immunized with 1 µg/100 µL of RBD-TT conjugates adjuvated with 500 µg of $Al(OH)_3$.

Rabbits (5 per group) received 3 µg/100 µL of RBD-TT conjugates adjuvated with 500 µg of $Al(OH)_3$ or 500 µg/100 µl of $Al(OH)_3$ as control.

40 hamsters were divided in two groups of 20 animals each. The first group received 3 µg/100 µL of the RBD-TT conjugate adjuvated with 500 µg of $Al(OH)_3$. The second group received 500 µg/100 µl of $Al(OH)_3$ as control.

**Anti-RBD IgG ELISA**

96 well ELISA plates (NUNC Maxisorp) were coated with 50 µL of RBD antigen at a concentration of 3 µg/mL with carbonate-bicarbonate buffer (pH 9.6) for 1 h at 37 ºC. Successively, plates were blocked in 5% skim milk-PBS for 1 h at 37 ºC. After 5 washes with PBS-0.5% Tween 20 (PBS-T), serum samples were added serially (1:3) diluted in PBS-1% BSA solution (pH 7.2) starting from 1/50. Plates were incubated for 1 h at 37 ºC and then washed again with PBS-T. Goat anti-mouse IgG-HRP antibody (Sigma Aldrich) diluted 1/5000 in PBS-1% BSA solution (pH 7.2) were added to each well and incubated for another hour at 37 ºC. Following 5 washes with PBS-T, 3',3'5,5'-tetramethylbenzidine (TMB) substrate was added to the plates and incubated for 20 minutes. Reactions were stopped with 2 N H2SO4 and the absorbance measured at 450nm in a microplate reader ELISA Multiskan EX (ThermoScientific). The endpoint titer was defined as the highest reciprocal dilution of serum that gives an absorbance 4-fold greater than pre-immune serum diluted 1/50.

**Molecular Virus Neutralization Assays**

The ability of anti-RBD specific antibodies to inhibit the RBD-ACE2 interaction was evaluated for a Molecular Virus Neutralization Assay. Briefly, microtiter plates (High binding, Costar, USA) were coated with 50 µL/well of ACE2-mFc (5 µg/mL) in carbonate-bicarbonate buffer, 0.1 M (pH 9.6) and incubated overnight at 4 °C. Plates were blocked with 200 µL/well of 2% of NFDM in PBST during 1h, at 37 °C. Serial dilutions of individual sera were mixed 1:1 (*v/v*) with RBD-hFc (40ng/mL) and incubated for 1h at 37 °C. Additional mixtures containing sera from mice of placebo groups were equally prepared, and considered as negative control. All samples were diluted in NFDM 0.2%/PBST (assay buffer). Mixtures were added to the plates (50 µL/well) and further incubated for 2 h at 37 °C. Next, 50 µL/well of alkaline phosphatase-conjugated anti-human IgG antibody (1:1000) were added followed by incubation for 1 h at 37 °C. Finally, 50 µL/well of *p*-nitrophenylphosphate (Sigma) diluted at 1 mg/mL in diethanolamine buffer (pH 9.8) were added, and plates were incubated at room temperature for 40 min. Absorbance at 405 nm was measured using a microwell system reader (Organon Teknica, Salzburg, Austria). All incubations were followed by three washing steps with PBST. Inhibition was calculated and expressed as percentage according to the formula: Inhibition (%)= [1-(A405 nm sample/A405 nm maximal recognition)]×100. Maximal recognition corresponds to RBD-hFc (40 ng/mL) mixed 1:1 (*v/v*) with assay buffer. For ID50 determination in the mVNA, dilutions were log transformed and data was adjusted to a log(inhibitor) *vs* normalized response with variable slope non-linear regression

**Virus neutralization assay**

The virus neutralization assay was performed following the recommendation of Manenti y cols with few modifications[6]. Animal serum samples were heat-inactivated for 30 minutes at 56 °C. Two-fold serial dilutions, starting from 1:10 to 1:2560 were then mixed with an equal volume of viral solution containing 100 TCID50 of SARS-CoV-2 (Strain 2025, Cuban Collection at National Laboratory of Civil Defense). The serum-virus mixture was incubated for 1 hour at 37 °C in a humidified atmosphere with 5% CO2. After incubation, 100 µL of the mixture at each dilution was added in duplicate to a cell plate containing a semiconfluent VERO E6 monolayer ($10^4$ cell/well). The plates were incubated for 4 days at 37 °C in a humidified atmosphere with 5% $CO_2$.

[6] Manenti A. Evaluation of SARS-CoV-2 neutralizing antibodies using a CPE-based colorimetric live virus micro-neutralization assay in human serum samples. *J Med Virol.* **2020**; 1–9.

After 3 days of incubation, the supernatant of each plate was carefully discarded and 100 µL of a sterile PBS solution containing 0.02% neutral red (Sigma, St. Louis, MO) was added to each well of the microneutralization plates. After 1 hour of incubation at room temperature, the neutral red solution was discarded and the cell monolayer was washed twice with sterile PBS containing 0.05% Tween 20. After the second incubation, the PBS was carefully removed from each well; then, 100 µL of a lysis solution made up of 50 parts of absolute ethanol (Sigma, St. Louis, MO), 49 parts of MilliQ and 1 part of glacial acetic acid (Sigma) was added to each well. Plates were incubated for 15 minutes at room temperature and then read by a spectrophotometer at 540 nm. The highest serum dilution, showing an optical density (OD) representing the 50% of average DO value from control cell wells, was considered as the neutralization titer and is represented as neutralizing titer 50 (NT50). Control cell wells involve VERO E6 monolayer with mixture of virus-serum. In case of serum sample where NT50 cannot be calculated until 1:2560 dilution, the assay was repeated starting with higher dilution.

**Detection of RBD-specific T cells by flow cytometry**

The RBD-specific memory T cells induced by RBD-TT conjugates (1 µg/100 µL of RBD-TT adjuvated with 500 µg of $Al(OH)_3$) was determined by flow cytometry. Splenocytes from immunized or control mice were added to 24 well plates at a density of 3×106/well and cultured for 72 h in RPMI medium 1640 (Gibco) supplemented with 10% (*v/v*) FBS (Capricorn), 100 U/mL penicillin-streptomycin, 1 mM pyruvate (Gibco), 50 µM β-mercaptoethanol and 50 UI/ml IL-2 (Sigma-Aldrich). Mouse splenocytes were stimulated with 5µg/mL of RBD and positive control wells were stimulated with 5 µg/mL ConA (Sigma-Aldrich) for 24 h. Brefeldin A (10 µg/mL; BD Biosciences) was administrated 6 h before staining. Then, the cells were harvested and stained first with a live/dead fixable near IR fluorescent dye (Invitrogen, by Thermo scientific) and the anti-CD4 PerCP/Cy5.5 (RM4-5), anti-CD8 PE (S3-6.7) and anti-CD44 PE-cy7 (IM7) surface markers (all from eBiosciences). Cells were subsequently fixed and permeabilized (eBiosciences) to staining with anti-IFNγ FITC (XM61.2) and anti-IL-4 APC (11B11) (all from eBiosciences). Flow cytometry data was acquired on a Gallios Cytometer (Beckman Coulter) and analyzed using Kaluza Software 1.2 version. For the gating strategy, CD4+ or CD8+ with CD44+ lymphocytes (defined as memory reactive RBD cells) were gated from live cells and the percentage of IFN-γ+ or IL-4+ were recorded.

**Quantification of cytokines by ELISA**

Supernatants from RBD-stimulated splenocytes were collected from 72-hour stimulation cultures and stored at -80 °C. Cytokines in cell culture supernatants were quantified using an IFN-γ and IL-4 ELISA kit (Mabtech) according to the manufacturer's instruction.

**Statistical analysis.** The analyses were performed with GraphPad Prism Software v.7.0.4. Two-sided nonparametric Mann–Whitney tests were conducted to compare differences between two experimental groups, Kruskal–Wallis ANOVA with Dunn's multiple comparisons tests were applied to compare >2 experimental groups. Antibody titer data were log transformed before analysis. P-values of $< 0.05$ were considered significant.

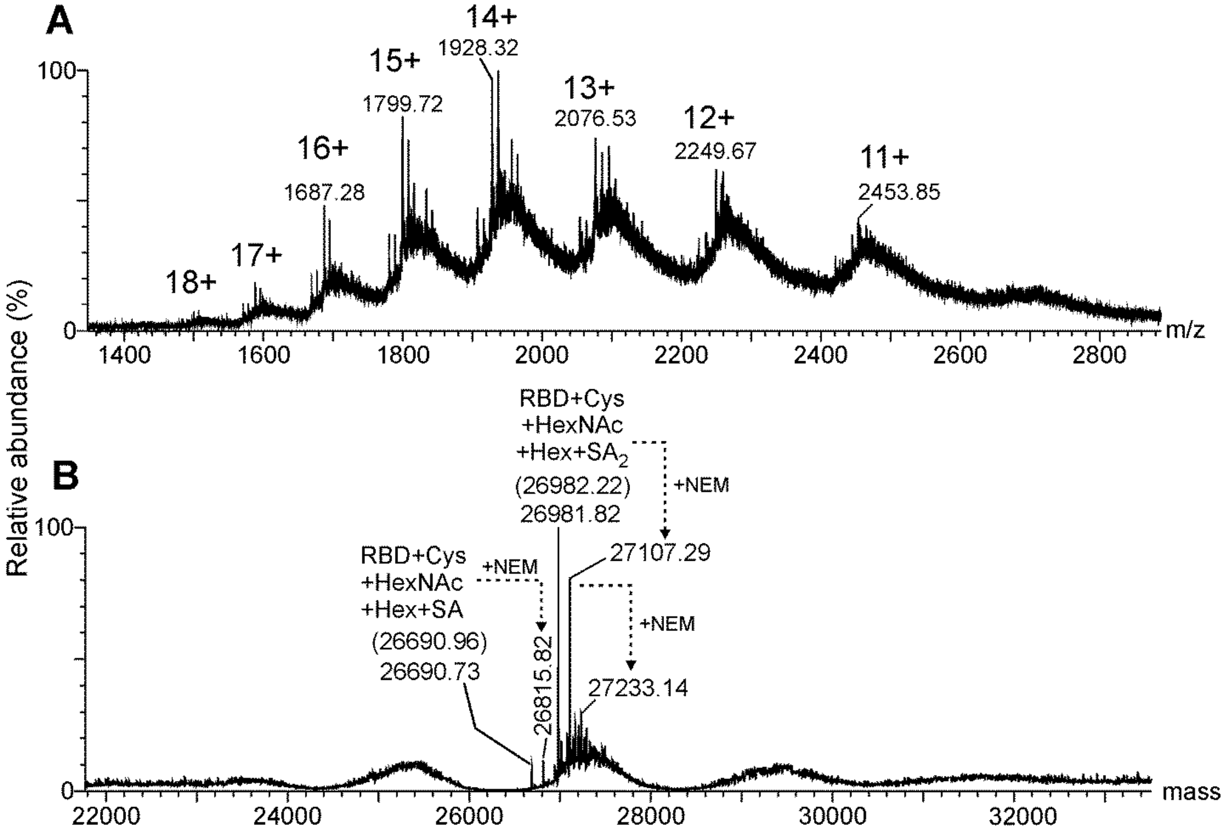


**Fig. S1.** (A) Multiply-charged and (B) deconvoluted ESI-MS spectrum of recombinant *N*-deglycosylated RBD Arg[319]-Phe[541]-(His)$_6$ with a theoretical molecular mass value of 26571.81 Da and 26863.07 Da, considering Cys[538] free and O-glycosylation (HexNAc-Hex-SA and HexNAc-Hex-SA$_2$, HexNAc: N-acetylhexosamine, Hex: hexose, SA: sialic acid). The signals with molecular masses of 26690.73 Da and 26981.82 Da, approximately 119 Da higher than expected, indicates that the protein is predominantly cysteinylated. Parentheses indicate the expected molecular masses for cysteinylated RBD and O-glycosylated. NEM: N-ethyl maleimide.

**Table S1.** Summary of the average molecular weight (MW) determined by ESI-MS of the RBD N-deglycosylated with PNGase-F and treated with N-ethylmaleimide.

| **Average M.W Theor. (Da) + 4 S-S + Cys[538]-SH + PNGase-F ($N^{331}$, $N^{343}$→D)** | **Modification [(a)]** | **Average M.W Theor. (Da)** | **Average M.W Exp. (Da)** | **Error (%) [(b)]** | **Assignment** |
|---|---|---|---|---|---|
| 25915.22 | + O-HexNAc-Hex-SA (+656 Da)<br>$C^{538}$-Cys (+119 Da) | 26690.96 | 26690.73 | 0.0009 | Monomer of RBD cysteinylated and O-glycosylated |
| | + O-HexNAc-Hex-SA (+656 Da)<br>$C^{538}$-Cys (+119 Da)<br>+NEM (+125 Da) | 26816.09 | 26815.82 | 0.001 | |
| | + O-HexNAc-Hex-$SA_2$ (+947 Da)<br>$C^{538}$-Cys (+119 Da) | 26982.22 | 26981.82 | 0.001 | |
| | + O-HexNAc-Hex-$SA_2$ (+947 Da)<br>$C^{538}$-Cys (+119 Da)<br>+NEM (+125 Da) | 27107.34 | 27107.29 | 0.0002 | |
| | + O-HexNAc-Hex-$SA_2$ (+947 Da)<br>$C^{538}$-Cys (+119 Da)<br>+ 2NEM (+250 Da) | 27232.47 | 27233.14 | 0.002 | |

(a) HexNAc: N-acetylhexosamine, Hex: hexose, SA: sialic acid, NEM: N-ethylmaleimide, S-S: disulfide bond
(b) Relative error (%) = | (M.W theor – M.W exp) / M.W theor | * 100.

.

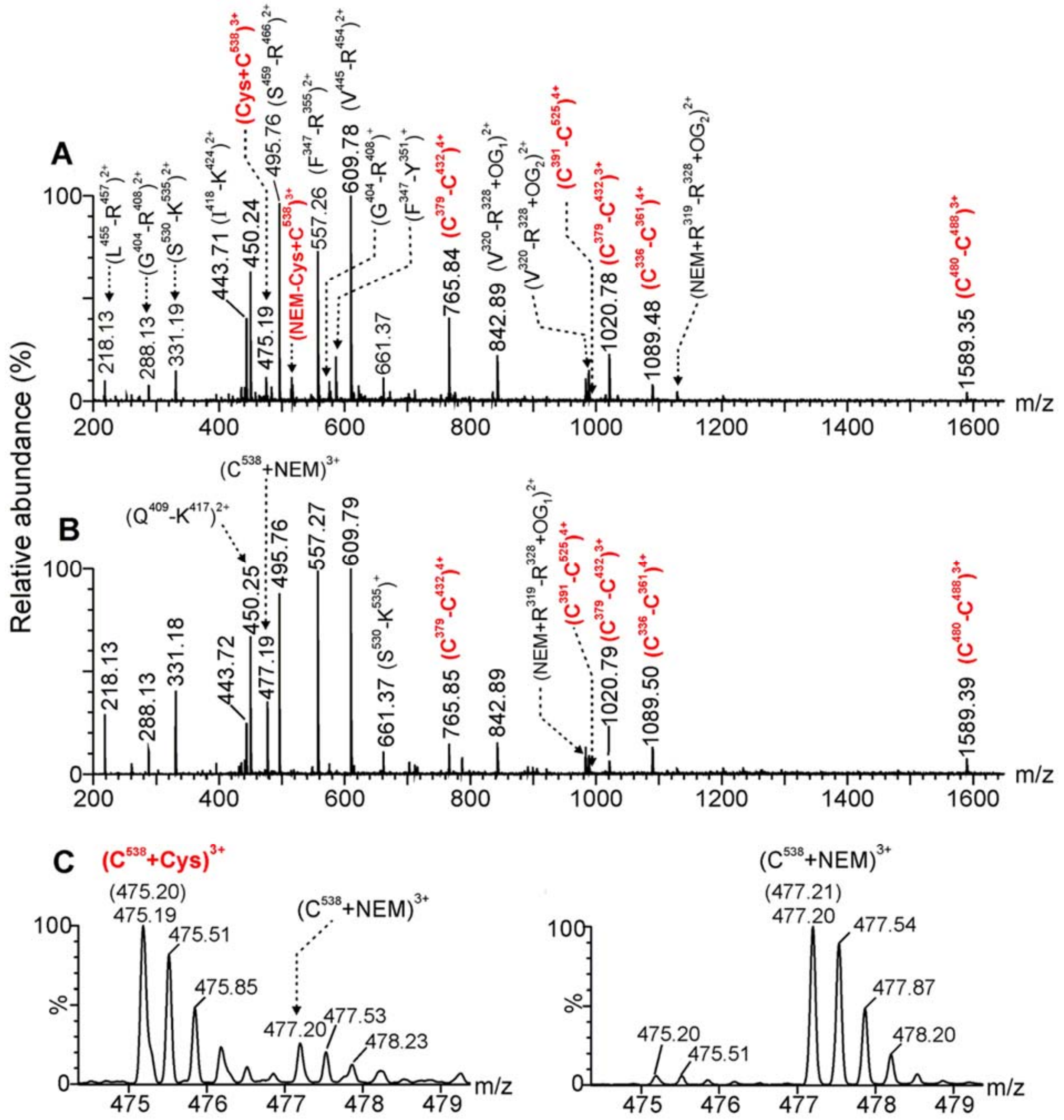


**Fig. S2.** (A) ESI-MS spectrum of the proteolitic peptides derived from RBD treated with NEM, deglycosylated with PNGase F and digested with trypsin. (B) ESI-MS spectrum of the peptides after the same protocol of (A), but with the previous reduction of the RBD with TCEP. (C) Expansion of the spectra at the region of the C-terminal peptide $C^{538}VNFHHHHHH^{547}$, confirming that the RBD was mostly cysteinylated at Cys538 (left panel, *m/z*: 475.20), while RBD reduction with TCEP set Cys538 free and then capped with NEM (right panel, *m/z*: 477.20).

**Table S2.** Sequence assignment for the tryptic peptides of the RBD Arg319-Phe541-(His)$_6$ and the treatment with TCEP generated by a Buffer-Free-Digestion protocol and analyzed by ESI-MS.

| Sequence assignment [b)] | *m/z* Exp RBD | *m/z* Exp RBD+TCEP | *m/z* Theor | z |
|---|---|---|---|---|
| $^{319}$RVQPTESIVR$^{328}$ | 592.84 | 592.84 | 592.84 | 2+ |
| NEM-$^{319}$RVQPTESIVR$^{328}$ | 655.36 | 655.34 | 655.37 | 2+ |
| NEM-$^{319}$RVQPTESIVR$^{328}$+*HexNAc+Hex+SA* (O-glycosylation) | 983.47 | 983.46 | 983.48 | 2+ |
| NEM-$^{319}$RVQPTESIVR$^{328}$+*HexNAc+Hex+SA$_2$* (O-glycosylation) | 1129.03 | 1129.03 | 1129.03 | 2+ |
| $^{319}$RVQPTESIVR$^{328}$+*HexNAc+Hex+SA* (O-glycosylation) | 920.96 | 920.95 | 920.96 | 2+ |
| $^{319}$RVQPTESIVR$^{328}$+*HexNAc+Hex+SA$_2$* (O-glycosylation) | 1066.50 | 1066.49 | 1066.50 | 2+ |
| $^{320}$VQPTESIVR$^{328}$ | 514.75 | 514.77 | 514.79 | 2+ |
| $^{320}$VQPTESIVR$^{328}$+*HexNAc+Hex+SA* (O-glycosylation) | 842.91 | 842.89 | 842.90 | 2+ |
| $^{320}$VQPTESIVR$^{328}$+*HexNAc+Hex+SA$_2$* (O-glycosylation) | 988.45 | 988.44 | 988.45 | 2+ |
| $^{329}$FPDITNL**C**PFGEVFDATR$^{346}$ **(C$^{336}$-C$^{361}$)**<br>$^{358}$ISN**C**VADYSVLYNSASFSTFK$^{378}$ | 1089.52<br>1452.37 | 1089.51<br>1452.34 | 1089.51<br>1452.35 | 4+<br>3+ |
| $^{347}$FASVYAWNR$^{355}$ | 557.28<br>1113.54 | 557.27<br>1113.54 | 557.28<br>1113.55 | 2+<br>1+ |
| $^{356}$KR$^{357}$ | 303.21 | 303.21 | 303.21 | 1+ |
| $^{379}$**C**YGVSPTK$^{386}$ **(C$^{379}$-C$^{432}$)**<br>$^{425}$LPDDFTG**C**VIAWNSNNLDSK$^{444}$ | 765.86<br>1020.81<br>1530.70 | 765.85<br>1020.82<br>1530.68 | 765.86<br>1020.81<br>1530.71 | 4+<br>3+<br>2+ |
| $^{387}$LNDL**C**FTNVYADSFVIR$^{403}$ **(C$^{391}$-C$^{525}$)**<br>$^{510}$VVVLSFELLHAPATV**C**GPK$^{528}$ | 992.53 | 992.52 | 992.52 | 4+ |
| $^{404}$GDEVR$^{408}$ | 575.27<br>288.13 | 575.26<br>288.14 | 575.28<br>288.14 | 1+<br>2+ |
| $^{409}$QIAPGQTGK$^{417}$ | 450.25<br>899.50 | 450.25<br>899.47 | 450.25<br>899.50 | 2+<br>1+ |
| $^{418}$IADYNYK$^{424}$ | 443.72<br>886.43 | 443.72<br>886.42 | 443.72<br>886.43 | 2+<br>1+ |
| $^{445}$VGGNYNYLYR$^{454}$ | 609.80<br>1218.58 | 609.79<br>1218.57 | 609.80<br>1218.59 | 2+<br>1+ |
| $^{455}$LFR$^{457}$ | 435.27<br>218.13 | 435.27<br>218.13 | 435.27<br>218.14 | 1+<br>2+ |
| $^{455}$LFRK$^{458}$ | 282.18 | 282.18 | 282.19 | 2+ |
| $^{458}$KSNLKPFER$^{466}$ | 373.54 | 373.54 | 373.55 | 3+ |
| $^{459}$SNLKPFER$^{466}$ | 495.77 | 495.76 | 495.77 | 2+ |
| $^{467}$DISTEIYQAGSTP**C**NGVEGFN**C**YFPLQSYGFQPTNGVGYQPYR$^{509}$<br>**(C$^{480}$-C$^{488}$)** | 1589.40<br>1192.31 | 1589.39<br>1192.29 | 1589.38<br>1192.29 | 3+<br>4+ |
| $^{529}$KSTNLVK$^{535}$ | 395.24 | 395.24 | 395.25 | 2+ |
| $^{530}$STNLVK$^{535}$ | 661.39<br>331.19 | 661.38<br>331.18 | 661.39<br>331.20 | 1+<br>2+ |
| $^{536}$NK$^{537}$ | 261.15 | 261.15 | 261.16 | 1+ |
| $^{538}$**C**VNFHHHHHH$^{547}$ (cysteinylated C$^{538}$) [a)]<br>**NH$_2$-C-COOH** | 475.19 | 475.20 | 475.19 | 3+ |
| $^{538}$**C**VNFHHHHHH$^{547}$ (cysteinylated C$^{538}$ + NEM-at the Nt) [a)]<br>**NEM-C-COOH** | 516.87 | 516.88 | 516.88 | 3+ |
| $^{538}$**C$_{NEM}$**VNFHHHHHH$^{547}$ (C$^{538}$ S-alkylated with NEM) [a)] | -<br>477.20 | 715.30<br>477.20 | 715.31<br>477.21 | 2+<br>3+ |

(a) *The six histidine residues located at the C-terminal end (residues 542-547) of the protein do not correspond to the RBD and were inserted in the cloning stage to facilitate the purification process of the recombinant protein by using IMAC.*

(b) *The superscript numbers indicate the location of the tryptic peptides within the analyzed protein. A brief description of the PTMs linked to the corresponding peptides is included. Nt means the N-terminal ends of the protein and the Cys residue linked by disulfide bond to Cys$^{538}$ were alkylated by a non-specific reaction with N-ethyl maleimide. HexNAc: N-acetyl hexosamine, Hex: hexose, NEM: N-ethyl maleimide, SA: sialic acid. C#-C# written in bold indicates the cysteine linked by native disulfide bonds.* ***C$_{NEM}$*** *indicates cysteine residues specifically alkylated with N-ethyl maleimide at the thiol group.* ***NH$_2$-C-COOH*** *corresponds to a cysteine residue linked by a disulfide bond to Cys$^{538}$. Nt indicates an N-terminal end. The residues indicated as D correspond to potential N-glycosylation sites located at Asn$^{331}$ and Asn$^{343}$ that were transformed into Asp by PNGase F.*

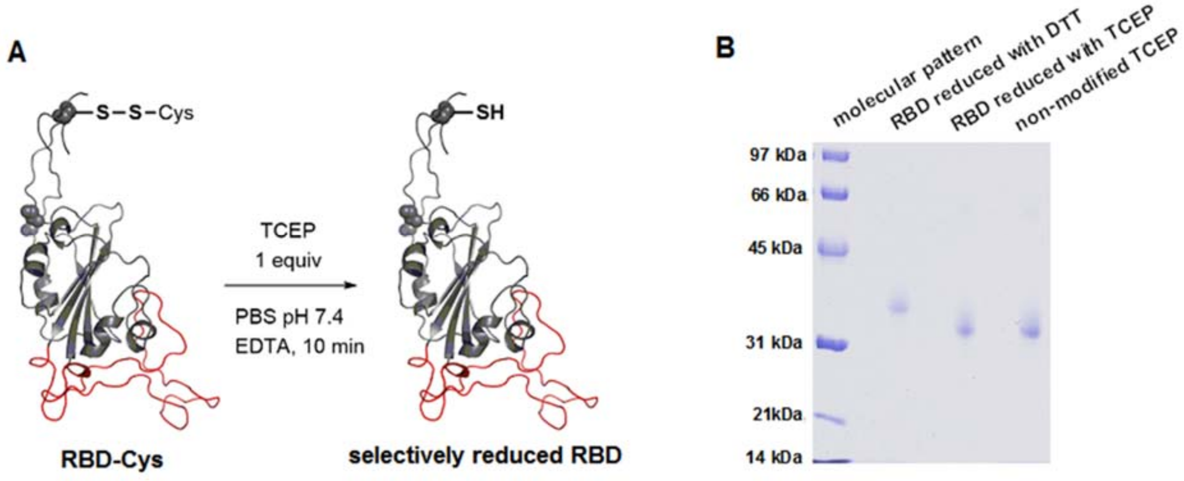


**Fig. S3.** (A) Reduction of recombinant RBD (Arg319-Phe541-(His)$_6$) with TCEP to selectively cleave the intermolecular disulfide bond between RBD's Cys538 and an additional Cys residue without affecting the intramolecular disulfide bridges. (B) SDS-PAGE analysis of the RBD reduced with DTT (proving the loss of the 3D structure) and TCEP in comparison with the non-modified recombinant protein.

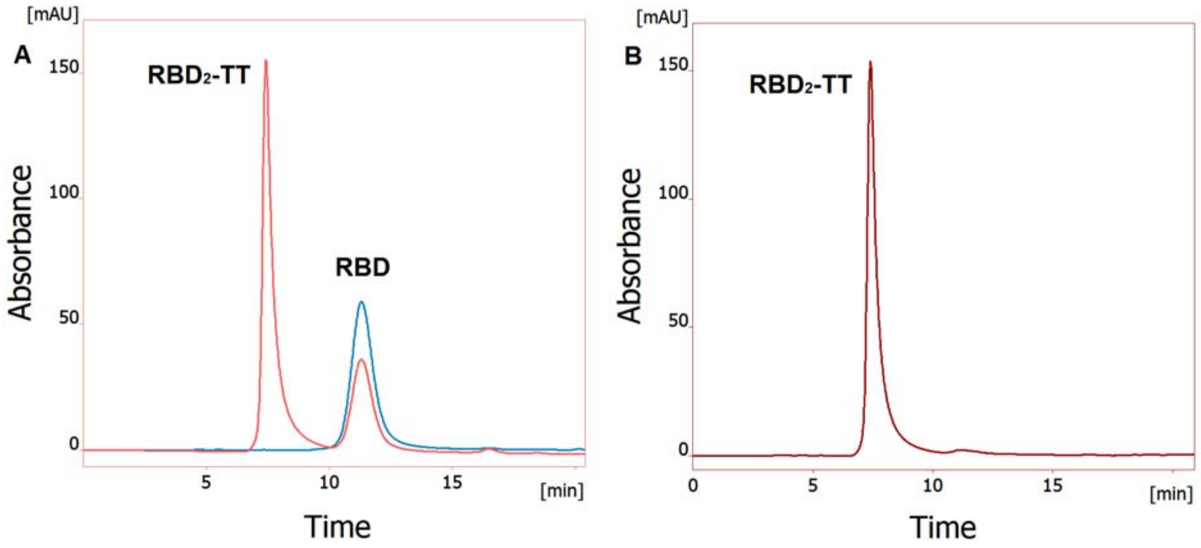


**Fig. S4.** SE-HPLC (Superdex 75®) chromatograms of (A) the crude conjugate $RBD_2$-TT overlapped with RBD and (B) purified $RBD_2$-TT.

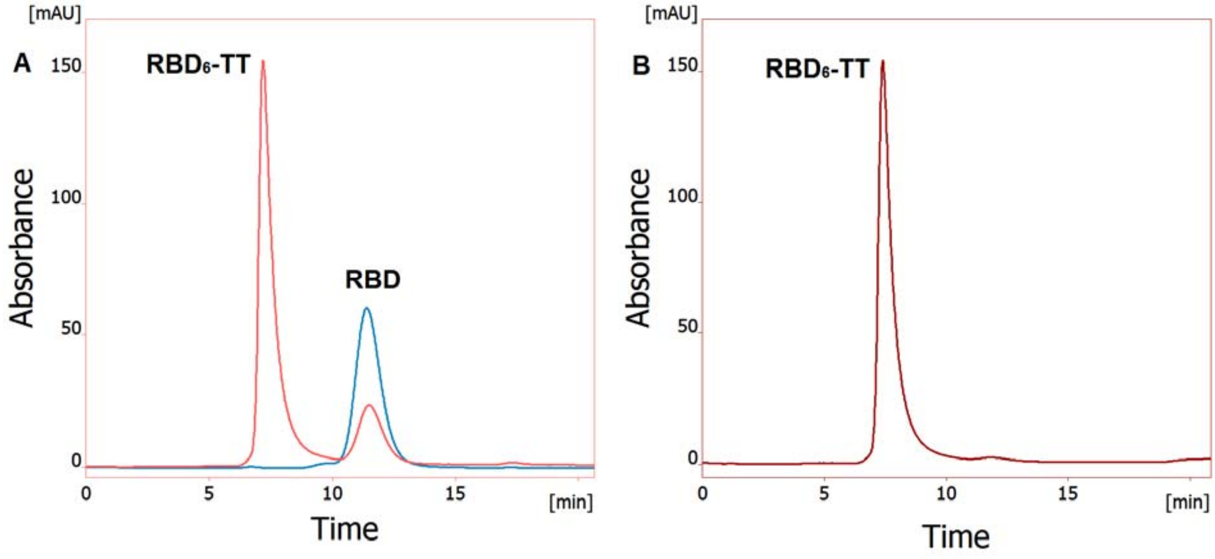


**Fig. S5.** SE-HPLC (Superdex 75®) chromatograms of (A) the crude conjugate $RBD_6$-TT overlapped with RBD and (B) purified $RBD_6$-TT.

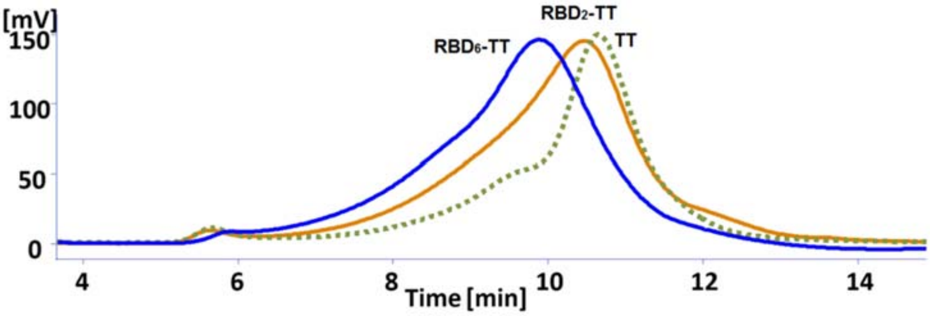


**Fig. S6.** Overlapped SE-HPLC (Superdex 200®) chromatograms of conjugates RBD6-TT (blue line), RBD2-TT (orange line) and TT (green dotted line).

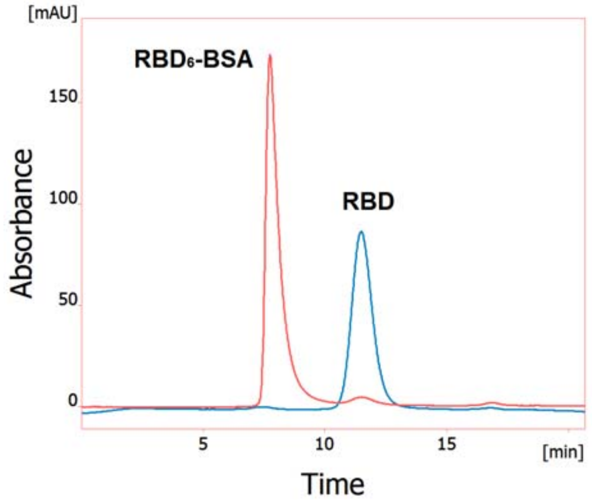


**Fig. S7.** Overlapped SE-HPLC (Superdex 75®) chromatograms of RBD6-BSA and RBD.

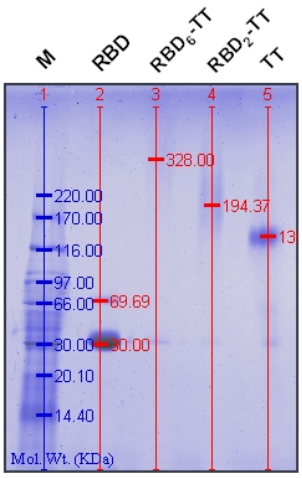


**Fig. S8.** SDS-PAGE analysis of the crude RDB2-TT and RDB6-TT conjugates in comparison with RBD and TT.

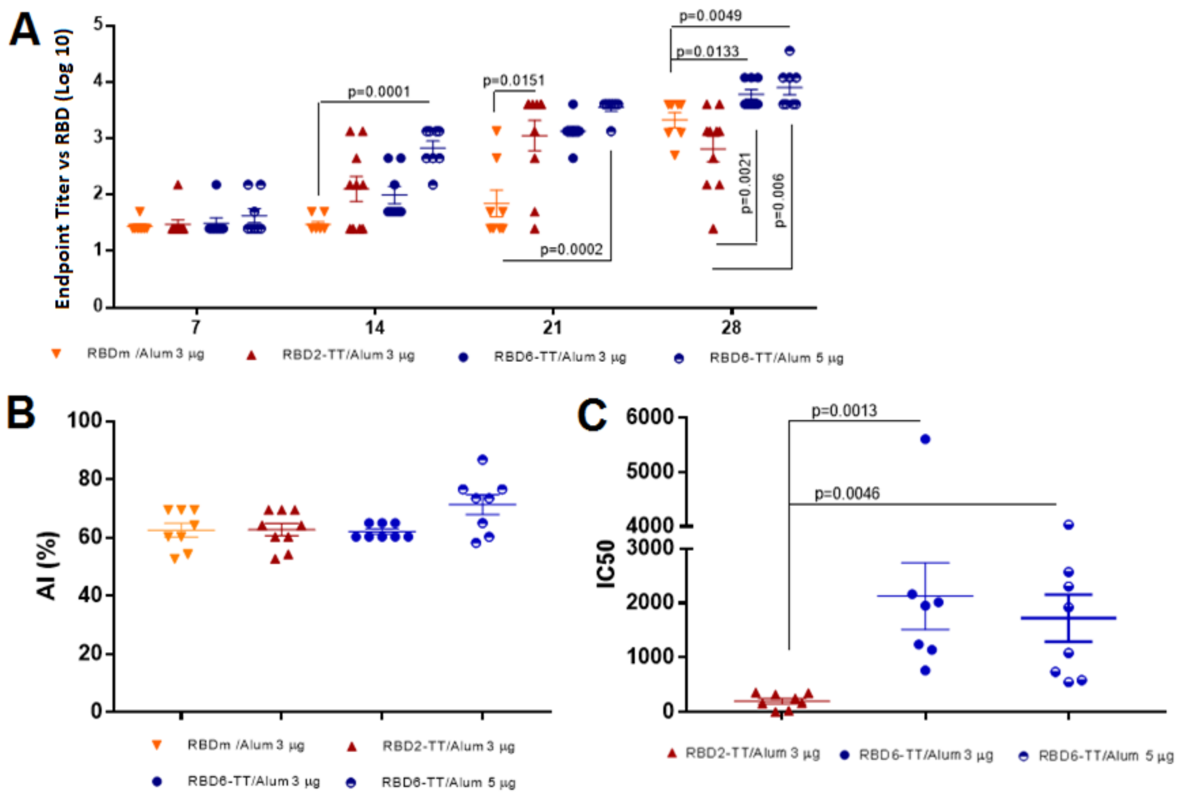


**Fig. S9.** Immune response in elderly C57BL/6 mice with 3µg of RBD$_2$-TT/Alum and 3µg or 5µg of RBD$_6$-TT/Alum compared to 3µg of monomeric RBD/Alum. (A) anti-RBD–specific IgG at days 7, 14, 21, and 28. (B) Avidity index of antibodies elicited by the four immunogens at T28. (C) Inhibition of the RBD-ACE2 interaction by antibodies elicited at T28.
